# Crosstalk between CD8^+^ T cells and systemic bile acid metabolism controls LCMV-induced immunopathology

**DOI:** 10.1101/2025.08.17.670599

**Authors:** Zsofia Keszei, Felix C. Richter, Henrique G. Colaço, Maximilian Baumgartner, Laura Antonio-Herrera, Magdalena Siller, Anna Hofmann, Csilla Viczenczova, Hatoon Baazim, Claudia D. Fuchs, Oleksandr Petrenko, Fabian Amman, Jakob-Wendelin Genger, Clarissa Campbell, Hanns-Ulrich Marschall, Thomas Reiberger, Michael Trauner, Andreas Bergthaler

## Abstract

Antiviral immunity has a profound effect on host metabolism, which can, in turn, modulate immune responses and influences disease pathology. Among its many functions, the liver orchestrates systemic bile acid (BA) metabolism, a pathway disrupted in chronic liver diseases such as viral hepatitis. BAs have become increasingly recognized for their immunomodulatory properties, and multiple BA species are being explored as therapeutic agents in liver diseases. Understanding the interplay between immune responses and BA metabolism could unlock new therapeutic opportunities based on BA modulation. Using lymphocytic choriomeningitis virus (LCMV) as a model, we investigated the interplay between chronic hepatotropic virus infection, BA metabolism and immunity. Our findings reveal that chronic LCMV infection increases BA levels and shifts circulating and liver BA composition towards host-derived, conjugated BAs. At the same time, hepatic BA transport and synthesis genes are broadly downregulated, which is at least partially dependent on CD8^+^ T cells. Additionally, we found that sustained high BA levels impact CD8^+^ T cell responses to chronic LCMV infection. Mice with elevated circulating BAs due to the lack of BA transporters OATP1a and OATP1b, showed impaired T cell expansion and reduced liver immunopathology. These findings reveal a reciprocal interplay between CD8^+^ T cells and BA metabolism, expanding our understanding of adaptive immunity against viral hepatitis. Moreover, it highlights how immuno-metabolic changes in liver disease may affect the body’s ability to fight infections and cancer.

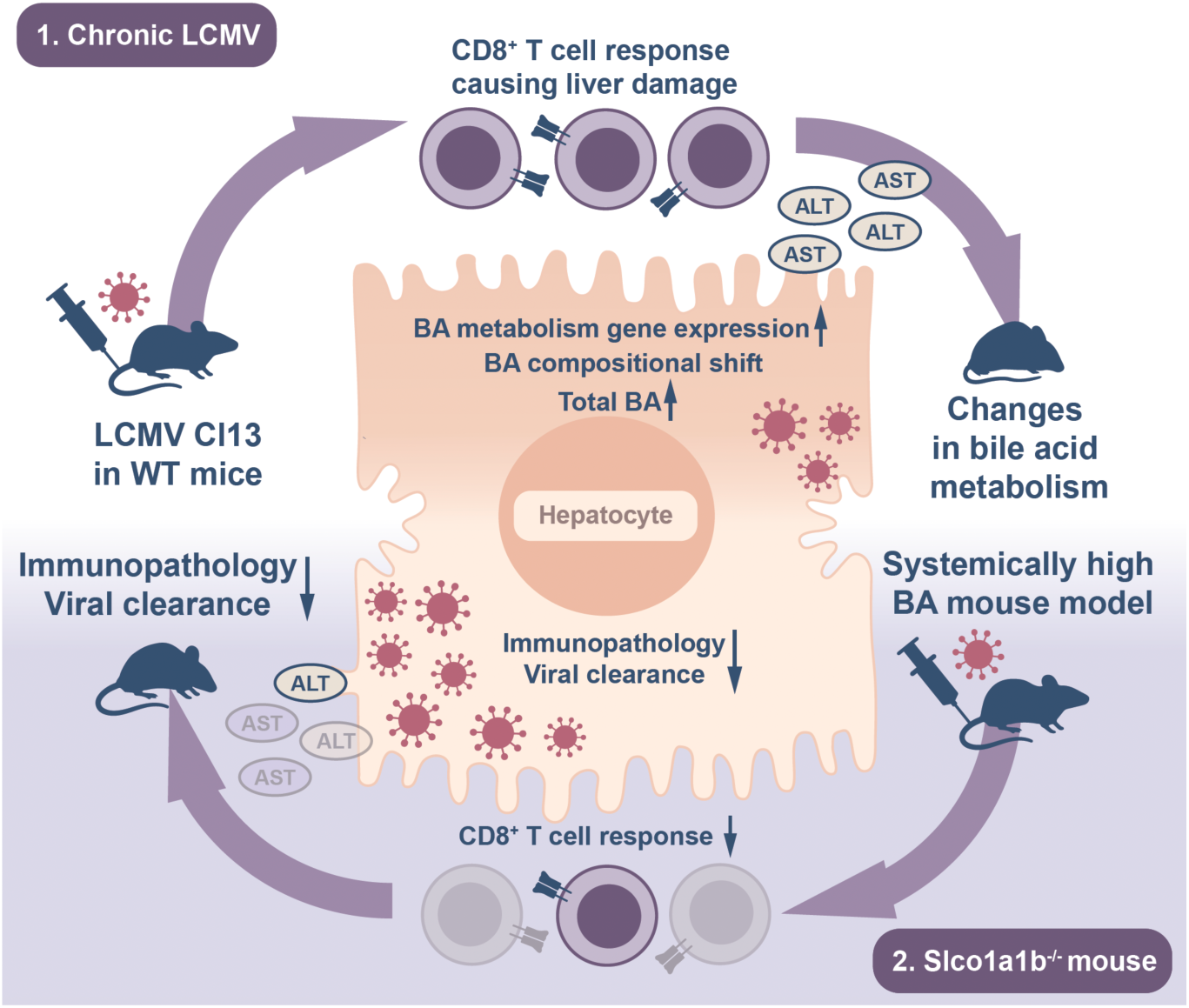

## Introduction

Responsible for two million deaths annually, liver disease accounts for 4% of all global deaths and ranks as the eleventh-leading cause of mortality worldwide. Viral hepatitis still accounts for most of these cases, followed by alcoholic liver disease (ALD) and metabolic dysfunction-associated steatotic liver disease (MASLD, formerly known as NAFLD), both of which are expected to increase worldwide (1).

The liver is a critical metabolic hub, also regulating systemic immunity against a variety of pathogens (2,3). One way it can influence systemic immunity is by modulating the levels of local and circulating metabolites (4,5), including bile acid (BA) species with immunometabolic properties (6). BAs are amphipathic cholesterol derivatives, which circulate between the liver and the intestine. In the liver, the steroid nucleus is conjugated with taurine or glycine, which makes them impermeable to cell membranes, reducing their toxicity and increasing their solubility. This enhanced solubility promotes the formation of micelles with lipids, facilitating lipid absorption in the intestine (7). Conjugated BAs are transported to bile canaliculi by the bile acid export pump (BSEP) encoded by the *Abcb11* gene. Then they travel through the gall bladder to the duodenum upon food uptake. Host-derived (also known as primary) BAs can go through extensive modifications by the gut microbiota to produce microbe-derived (also known as secondary) BAs. These changes start with deconjugation catalyzed by bacterial bile salt hydrolases (BSH), a necessary step preceding subsequent modifications (8). Over 95% of BAs are reabsorbed by ileal enterocytes and then enter the portal vein blood flow back to the liver (9). In the liver, BA uptake is mediated by basolateral transporters, including the Na^+^-taurocholate co-transporting polypeptide (NTCP) encoded by *Slc10a1*, and the organic anion-transporting polypeptides (OATPs), specifically OATP1b2 (*Slco1b2*), OATP1a1 (*Slco1a1*), and OATP1a4 (*Slco1a4*) in mice (10). While there is considerable overlap in substrate specificity, NTCP and OATPs each play a critical role in the uptake of conjugated and unconjugated BAs, respectively (11–13). Genetic ablation of either NTCP/*Slc10a1* or OATPs result in altered BA pools, demonstrating the role of these transporters in influencing the serum levels of different BA species (11,14–16). Moreover, suppression of BA uptake transporters has been described in liver inflammation, potentially protecting hepatocytes from excessive BA concentrations (17,18).

*De novo* BA synthesis in hepatocytes is regulated by a well-described negative feedback loop. BAs are transported into ileal enterocytes by the apical sodium-dependent bile acid transporter (ABST/*Slc10a2*), and induce fibroblast growth factor 15 (*Fgf15*) expression via activation of the farnesoid X receptor (FXR/*Nr1h4*). FGF15 secreted into portal circulation functions as a crucial hormone regulating BA metabolism in the liver by binding to fibroblast growth factor receptor 4 (FGFR4) (19). Through the activation of the transcriptional co-repressor Small Heterodimer partner (SHP/*Nr0b2*), FGFR4 signaling initiates the downregulation of the rate-limiting BA synthesis enzyme *Cyp7a1*. In addition, hepatic FXR activation can directly induce the expression of *Shp* (20,21). Several BA transporters were also reported to be regulated by FXR*/Nr1h4* or SHP*/Nr0b2*, contributing to hepatocellular homeostasis by limiting BA uptake. (22–26).

Liver diseases including viral hepatitis (27–31) are commonly associated with perturbation in BA metabolism resulting in altered BA composition (6). Such modulations of the BA pool can have a far-reaching impact on innate (32–34) and adaptive immune cell functions. Direct effects of BAs on adaptive immune cells have only recently been discovered. Multiple microbe-derived BA species modulate CD4^+^ T cell subsets by promoting intestinal regulatory T cell formation and function (35–37). In addition, microbial-derived BAs have immunomodulatory effects on CD8^+^ T cells by regulating both their metabolism and effector function through FXR or TCR signaling (38–41). Due to the pivotal role of CD8^+^ T cells in combating both intracellular pathogens and cancer, understanding their modulation by BAs is of crucial importance. However, it is not yet clear how endogenous BAs are modulated upon chronic viral infection and whether they, in turn, influence the antiviral immune response.

Thus, we set out to explore the link between liver diseases and BA dysregulation by exploiting the viral hepatitis model of LCMV Cl13. This benchmark model of viral persistence in mice (42,43) infects hepatocytes, among other cells, and triggers a CD8^+^ T-cell-mediated immunopathology (44–46). First, we assessed the impact of chronic viral infection on changes in BA metabolism and second, the effects of systemically high BA levels on cytotoxic T cell-mediated immunity and host response to liver injury. We found that LCMV infection substantially elevated systemic BA levels and altered the composition of circulating BA species in wild-type mice. In parallel, we observed a downregulation of hepatic BA metabolism that was at least partially dependent on CD8^+^ T cells. To assess the impact of systemically high BA levels, we employed a mouse model of genetic deletion of the hepatic BA transporters *Slco1a* and *Slco1b*. This model revealed that high levels of BAs reduced CD8^+^ T cell-mediated immunity and accordingly dampened liver immunopathology during chronic LCMV infection. Thus, our data demonstrate a novel reciprocal interplay between CD8^+^ T cells and systemic BA levels during hepatic injury.

## Results

### Chronic LCMV infection shifts BA levels and composition

To establish if BA metabolism is affected by viral infections, we quantified total BA species in the serum of mice infected with 2×10^6^ focus forming units of LCMV strain clone 13 (Cl13). We observed an increase in total serum BAs which was most pronounced 12 days post-infection (Figure 1a), confirming previous observations with another strain of LCMV (47). To validate our results and gain novel insights into the composition of BA levels during LCMV infection, we performed more sensitive analysis by targeted liquid chromatography-mass spectrometry (LC-MS) across several time points on liver and serum samples. Supporting our initial findings, we observed a profound alteration of BA profiles as the viral infection progressed (Figure 1b). This shift was marked by an enrichment of predominantly host-derived and conjugated BAs in both serum and liver, which was most pronounced 8 days post-infection (Supplementary figure 1a, b), which coincides with the peak of CD8^+^ T cell responses (48).

**Figure 1:**
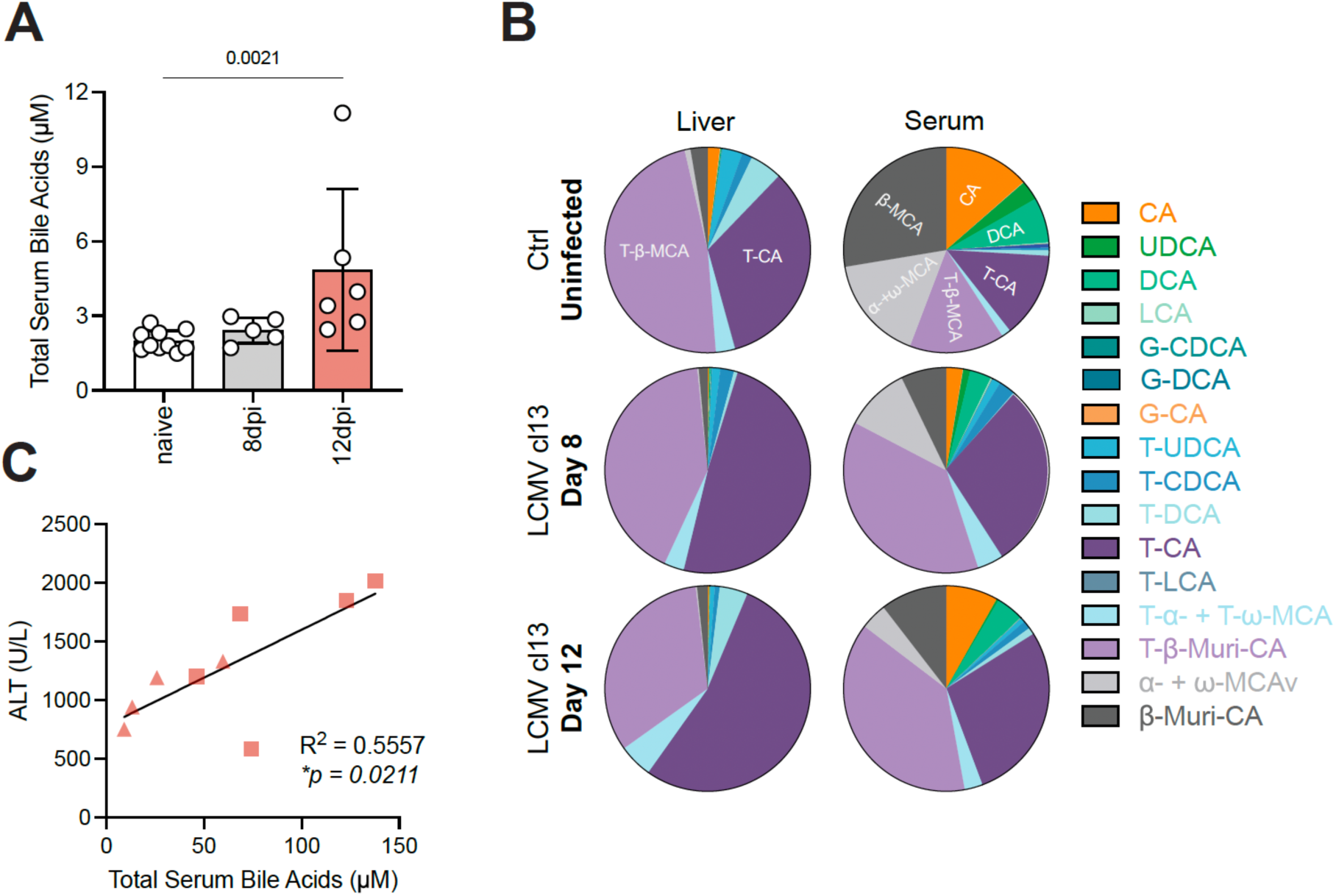
Viral hepatitis alters hepatic and systemic BA levels and composition. (a) Total serum BA levels of C57BL6/J mice infected with LCMV Cl13 quantified by colorimetric total BA assay. (b) Composition of serum and liver BA species in uninfected and LCMV Cl13-infected mice at 8- and 12-days post infection measured by LC-MS. (c) Correlation of hepatic damage marker ALT with systemic BA levels in the serum of C57BL6/J mice infected with LCMV Cl13 at day 12 post-infection measured by LC-MS. Data presented as mean ± SEM. Data from one experiment (n = 6-10 mice/group) (a). Pooled data from two independent experiments (n = 4-5 mice/experiment) (a,b). Shapes indicate each experiment (c). Kruskal Wallis test (a). Simple linear regression (c).

Host-derived conjugated BAs are deconjugated by microbial BA hydrolases (BSH) in the lower gastrointestinal tract. This process is pivotal for subsequent microbial modifications (8,49). During LCMV infection, the gut microbiota undergoes a substantial compositional shift, causing gut dysbiosis (50). To examine whether the shift towards host-derived conjugated BA species may be caused by a decrease of BSH-expressing gut microbes, we performed a PICRUSt analysis (51) on published 16S rRNA sequencing data from the colon of LCMV-infected mice 8 days post-infection (50). Surprisingly, the results showed a relative increase of BSH-expressing bacteria upon viral infection. (Supplementary figure 1c). This suggests that the accumulation of host-derived conjugated BAs in the serum and liver are likely not driven by an inability of the microbiota to process these BAs.

Alternatively, it seemed likely that this accumulation could result from a direct effect of liver damage (18). Indeed, there was a positive correlation between the liver damage marker alanine aminotransferase (ALT) and the total concentration of BAs at day 12 post-LCMV infection as quantified by LC-MS BA profiling (Figure 1c). Thus, direct release of host-derived conjugated BAs into the bloodstream due to hepatocyte damage could explain the observed shift in circulating BA profiles.

### Chronic LCMV infection decreases hepatic BA metabolism gene expression

In addition to the systemic changes in BA levels and composition, we wanted to assess the impact of LCMV infection on organismal BA metabolism. In the liver, we observed that BA transport and synthesis was substantially altered by LCMV infection on both transcriptomic and proteomic level. KEGG pathway enrichment analysis of a published dataset (52) revealed a broad metabolic reprogramming in the liver during viral infection, with numerous metabolism-related KEGG pathways downregulated especially on day 8 post-infection (Supplementary figure 2a), while ample immune-related pathways were upregulated as expected (Supplementary figure 2b). Bile secretion was among the most downregulated KEGG pathways, together with other terms related to fatty acid- and steroid metabolism (Supplementary figure 2a). It included genes responsible for primary BA synthesis, basolateral BA transport from the blood to the hepatocyte, as well as bile secretion, as shown on the schematic (Figure 2a). Transcripts of basolateral BA uptake transporter genes (*Slc10a1*, *Slco1a1*, *Slco1a4* and *Slco1b2*), canalicular BA export pump *Abcb11* and BA biosynthesis (*Cyp7a1, Cyp27a1, Cyp39a1*) were all found downregulated at least in one timepoint across both transcript and protein levels (Figure 2b).

**Figure 2:**
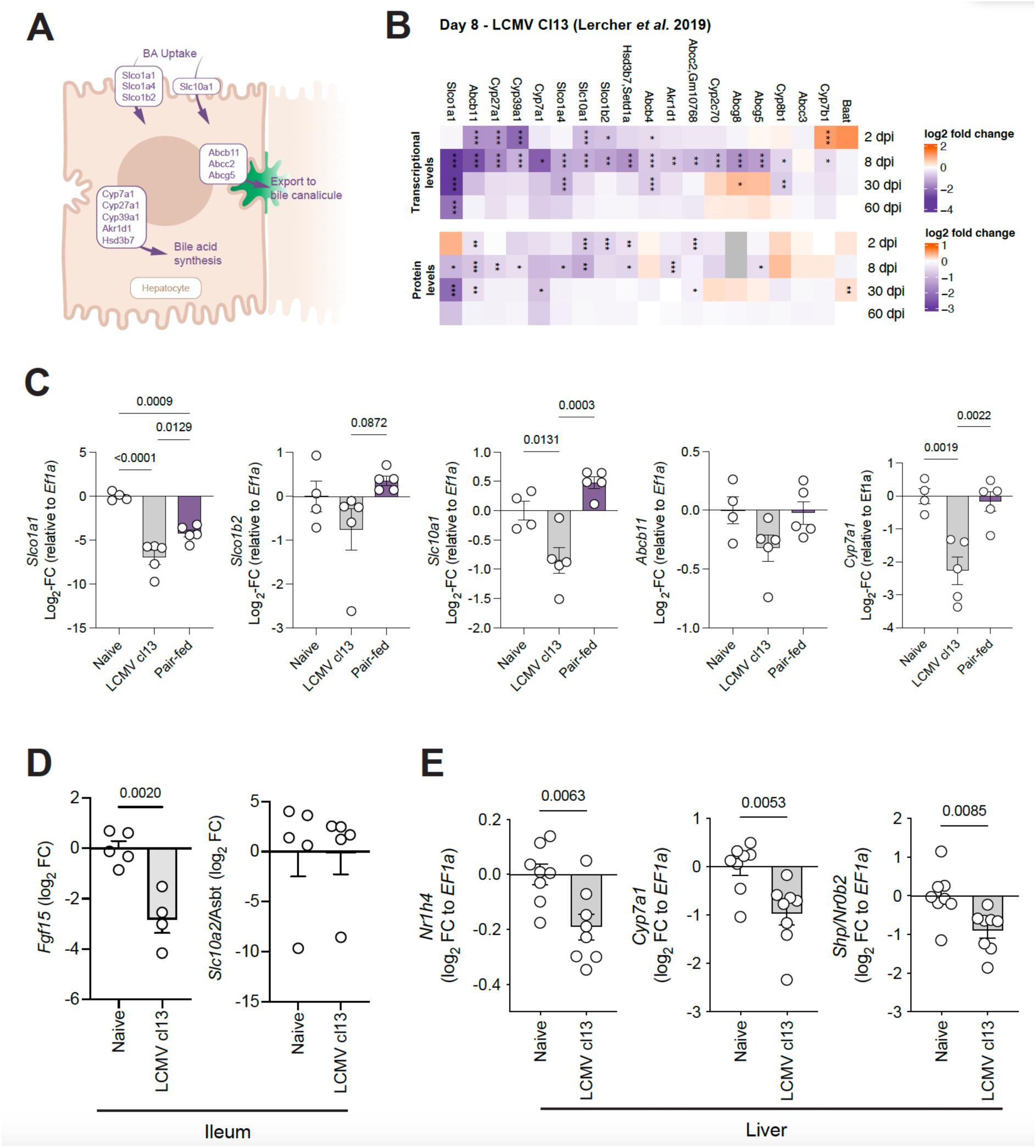
Chronic LCMV infection decreases the expression of genes involved in hepatic BA regulation, synthesis and transportion. (a) Overview schematic of BA homeostasis-related genes in hepatocytes that are significantly downregulated at both the transcript and protein levels at one or more time points. (b) Expression of BA homeostasis-related genes and proteins during infection with LCMV Cl13 infection based on Lercher *et al*. (52). (c) Hepatic gene expression of BA metabolic genes in response to LCMV Cl13 infection or pair-feeding. (d) Gene expression in ileum of uninfected mice and LCMV Cl13-infected mice at day 8 post-infection. (e) Expression of hepatic genes involved in FGF15-SHP signaling. Data presented as mean ± SEM. Data from one experiment (n = 4-5 mice/group) (c,d) and pooled from two independent experiments (n = 8 mice/group) (e). (c) One-Way ANOVA. (d-e) Unpaired Student’s t-test.

We next sought to identify the driving factors behind these transcriptional changes. Chronic LCMV infection induces anorectic behavior (53), which can have significant impact on the regulation of BA metabolism and associated gene expression (19,54). To confirm that the alterations in gene expression are not solely due to reduced food intake, we evaluated the expression of BA transporters and rate-limiting BA synthesis enzyme *Cyp7a1* in uninfected mice that were fed an equivalent amount of food as their Cl13-infected counterparts (pair-fed mice). We observed similar expression levels in both uninfected and pair-fed mice (Figure 2c), indicating that the alterations in transcript levels are not solely influenced by reduced food intake, but likely require an inflammatory response.

The downregulation of BA metabolism in hepatocytes during chronic LCMV infection could be caused by increased FGF15 signaling *via* the FGFR4 (*Fgfr4*)-SHP (*Nr0b2*) axis (19). Alternatively, hepatic FXR (*Nr1h4*) can induce SHP (*Nr0b2*) expression (20,21) or directly regulate BA transport and synthesis genes (22–26). To exclude that ileal FGF15 may be involved in the regulation of hepatic BA metabolism in our setting, we measured the expression of key target genes in both the ileum and liver. Ileal *Fgf15* expression was reduced, while we observed no difference in the expression of the ileal uptake transporter ASBT/*Slc10a2* (Supplementary figure 2c) suggesting that the regulation of hepatic BA synthesis and transport genes may be independent of FGF15 signaling. In line with this, we observed that transcripts encoding for FXR (*Nr1h4*), SHP (*Nr0b2*) and the FXR-target gene *Cyp7a1* were all reduced in the liver of Cl13 infected mice at 8 days post-infection relative to naïve animals (Figure 2d). Together, our data suggest that the downregulation of BA metabolism-related genes in the liver may be independent of ileal FGF15 signaling and reduced food intake. Taking into account the correlation of liver damage with circulating BA levels, we propose that the observed perturbation may be a direct consequence of the immune activation on liver gene expression.

### CD8^+^ T cells play an important role in downregulating BA transporter expression

Infection with LCMV is characterized by an early type-I interferon (IFN-I) response, a potent innate anti-viral cytokine, which can drive substantial metabolic alterations in the liver (52). To test whether IFN-I signaling controls BA transporter expression during viral infection, we genetically ablated the Ifn receptor alpha (*Ifnar*) in the whole mouse (*Ifnar1^-/-^*) or specifically in the liver (*Alb-Cre x Ifnar1^fl/fl^*). Upon infection with LCMV Cl13, mice with either whole body or liver-specific *Ifnar*-deficiency exhibited similar reductions in hepatic BA transporter expression (Supplementary Figure 3a+b) compared to littermate controls. This suggests that the downregulation of BA transporters is independent of IFN-I signaling.

To address whether the adaptive immune response to Cl13 is required for the changes in BA-metabolic gene expression, we infected *Rag2^-/-^* mice which are unable to produce mature T and B lymphocytes (55) with LCMV Cl13. Interestingly, *Rag2^-/-^* mice were protected from infection-associated downregulation of all tested genes (Figure 3a). This implicates the adaptive immune response in regulating genes involved in BA transport and synthesis. To further pinpoint the key players involved, we sought to determine if CD8^+^ T cells drive the downregulation of these transporters. Administering CD8α-depleting antibodies successfully depleted CD8^+^ T cells from circulation (Supplementary Figure 3c) resulting in similar modulation of BA-metabolic gene expression as seen in *Rag2*^-/-^ mice, albeit with slightly attenuated outcomes (Figure 3b). CD8^+^ T cells can influence target cells through secretion of various inflammatory cytokines. To assess the impact of CD8^+^ T cell effector cytokines on BA transporter gene expression, we administered blocking antibodies against IL-6, TNF-α or IFN-γ every second day. None of these treatments prevented the downregulation of basolateral BA transporter expression upon LCMV infection (Supplementary figure 3d-f). Only the downregulation of *Abcb11*, which encodes the canalicular transporter BSEP, was significantly attenuated upon IL-6 and TNF-α blockade (Supplementary figure 3g), in line with previous reports indicating that TNF-α can regulate *Abcb11/*BSEP expression *in vitro* (56). Together, this data demonstrates the role of adaptive immunity, in particular CD8^+^ T cells, in the transcriptional downregulation of hepatic BA transporters during LCMV Cl13 infection. These results further support our hypothesis that immune activation drives alterations in BA levels and BA transporter expression, highlighting the central role of CD8^+^ T cells in these changes.

**Figure 3:**
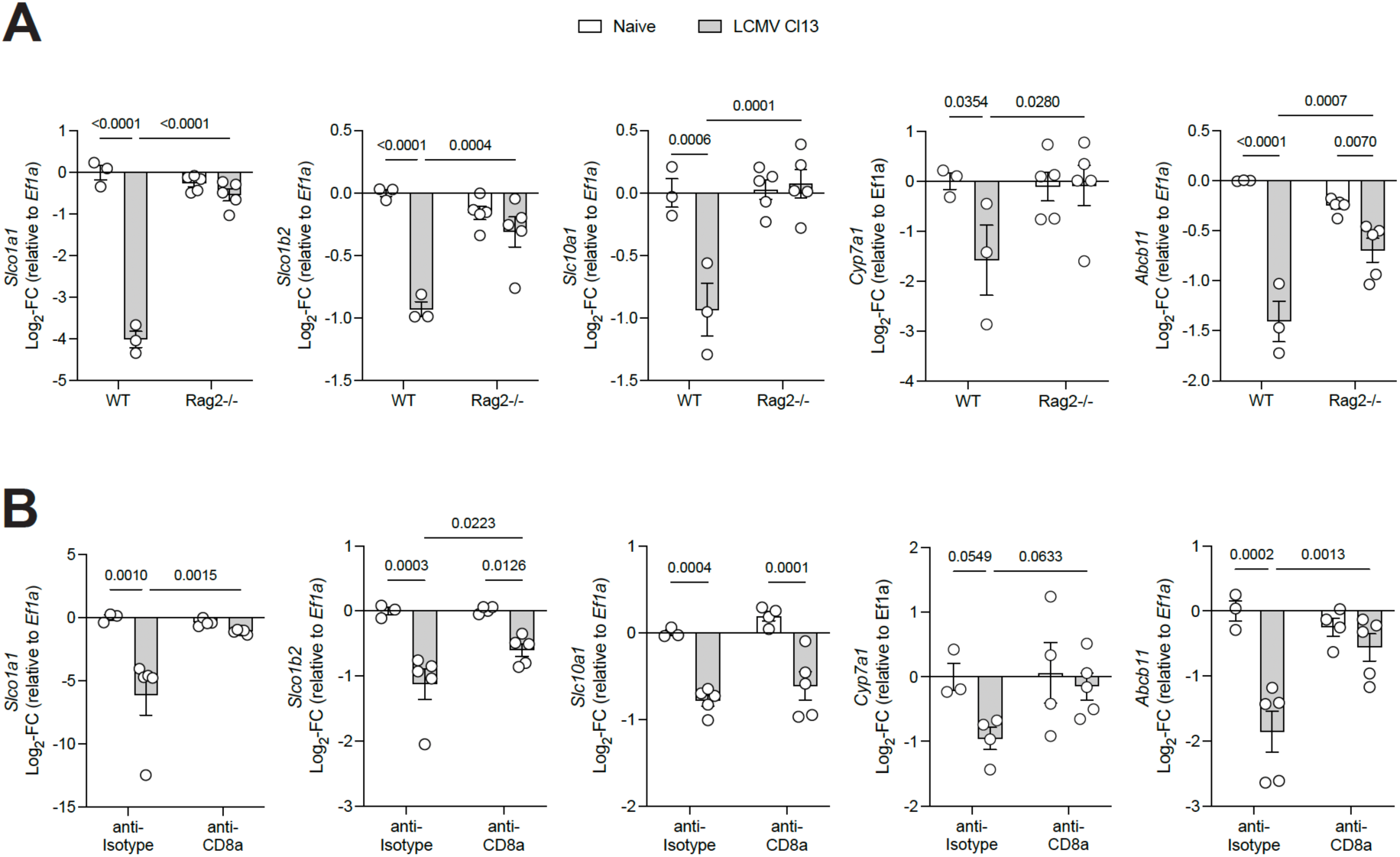
Downregulation of hepatic BA transporters are partially due to CD8^+^ T cells during LCMV Cl13 infection. (a) Hepatic expression of BA metabolism genes in response to LCMV Cl13 infection in C57BL6/J and *Rag2*-deficient mice. (b) Hepatic BA metabolism gene expression in response to LCMV Cl13 infection and CD8a-depleting antibody administration. Data presented as mean ± SEM. Representative data from two independent experiments. (a,b) Two-Way ANOVA.

### Splenic CD8^+^ T cell response is impaired in mice with genetic loss of BA transporters OATP1A/1B

Our results so far showed an increased abundance and changed composition of BAs during LCMV Cl13 infection. Considering the far-reaching immunomodulatory impact of BAs on CD8^+^ T cells (38,39,41), we next sought to understand the impact of high BA levels on the CD8^+^ T cell response during LCMV Cl13 infection. To address this, we made use of the mouse model *Slco1a1b^-/-^* (hereafter referred to as *Slco^-/-^*), which maintain systemically high BA levels due to lack of the entire locus containing the BA transporter genes *Slco1a* and *Slco1b* (11). First, we confirmed the *Slco^-/-^*genotype by verifying the deletion of genes involved in the cassette including *Slco1a1* (Supplementary figure 4a). As previously described (11), this deletion led to elevated levels of circulating total BAs (Supplementary Figure 4b) and bilirubin (Supplementary Figure 4c) in naïve mice. Upon infection of *Slco^-/-^* and littermate controls with LCMV (Figure 4a), we observed a decrease in virus-specific CD8^+^ T cells in the spleens of *Slco^-/-^* compared to littermate controls (Figure 4b+c), while total splenic CD8^+^ T cell numbers remained comparable (Supplementary Figure 4d). Interestingly, splenic CD8^+^ T cells in *Slco*^-/-^ mice exhibited reduced expression of the activation markers CD44 and PD1 (Figure 4d+e), indicating a disruption in T cell activation. In line with this, we found that the percentage of CD8^+^ T cells expressing the cellular proliferation marker KI67^+^ was reduced in *Slco^-/-^* mice (Supplementary Figure 4e).

**Figure 4:**
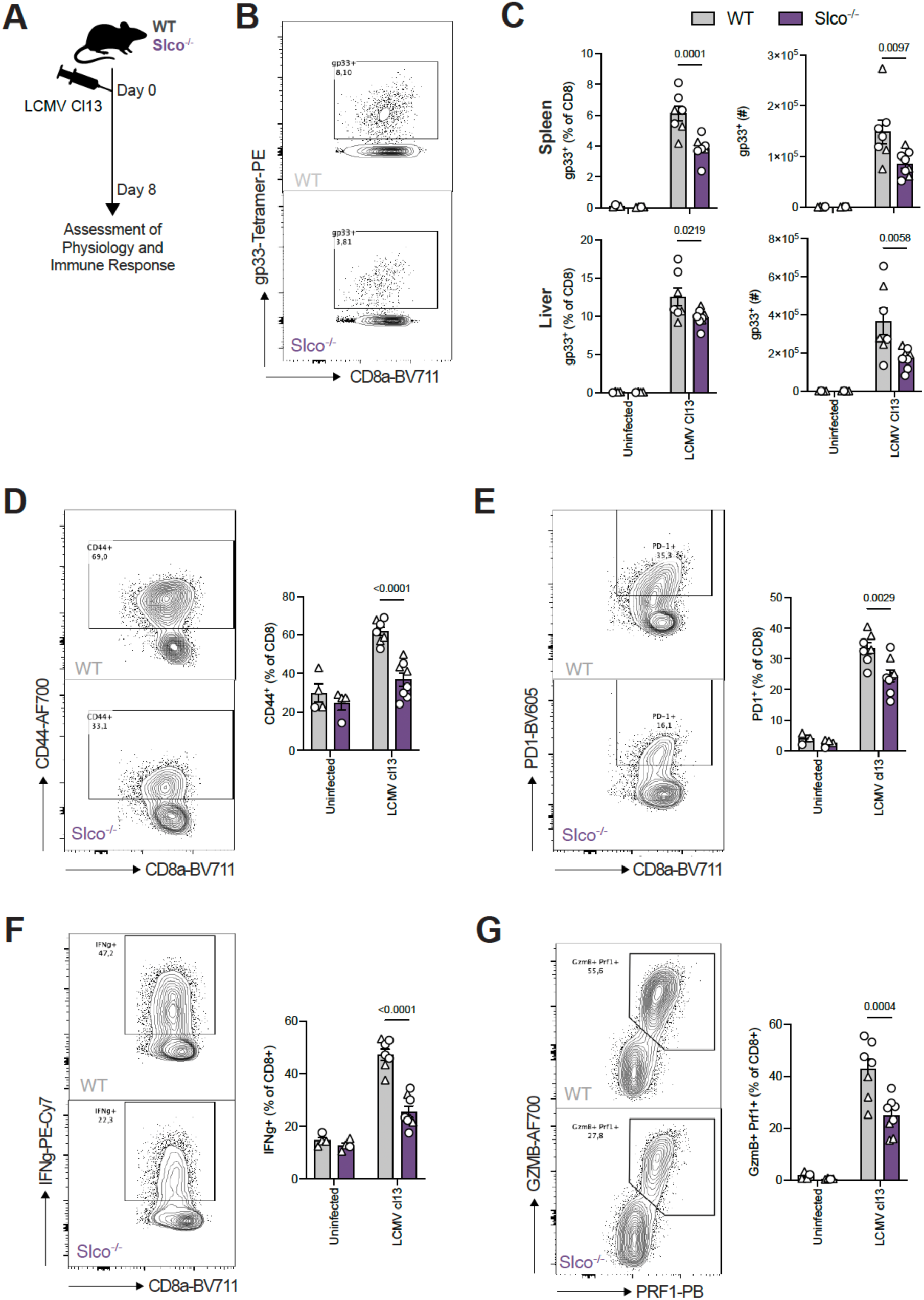
Impaired splenic CD8^+^ T cell response to LCMV Cl13 infection in mice with systemically elevated BAs. (a) Schematic of experimental design of LCMV infection in *Slco^-/-^* mice. (b) Representative flow cytometry plot of LCMV GP33-specific CD8^+^ T cell in the spleen 8 days post LCMV Cl13 infection. (c) Frequency and total number of LCMV GP33-specific CD8^+^ T cells in spleen and liver at day 8 post-infection. (d,e) Frequency of (d) CD44- and (e) PD1-expressing CD8^+^ T cells in spleen at day 8 post-LCMV Cl13 infection. (f) Frequency of IFN-γ-producing splenic CD8^+^ T cells after 4 hours restimulation with PMA and ionomycin at 8 days post-infection. (g) Frequency of GZMB^+^ PRF1^+^ splenic CD8^+^ T cells at day 8 post-LCMV Cl13 infection. Data presented as mean ± SEM. Data pooled from two independent experiments; each experiment represented in a different shape (n = 4-8 mice/group) (c-g) Two-Way ANOVA.

To test whether splenic CD8^+^ T cells themselves were less functional, we first restimulated total splenocytes with PMA/Ionomycin. Upon restimulation, the percentage of total CD8^+^ T cells producing key effector cytokine IFN-γ was drastically reduced (Figure 4f). Similarly, the percentage of CD8^+^ T cells producing the cytolytic proteins Granzyme B (GZMB) and Perforin (PRF1) was decreased (Figure 4g). These findings are in line with the reduced activation and presence of virus-specific CD8^+^ T cells in the spleen of *Slco^-/-^* mice. To assess whether virus-specific CD8^+^ T cells show any impaired function, we next restimulated isolated lymphocytes with the immunodominant viral epitope GP_33-41_. Interestingly, we found no differences in the production of effector molecules for either IFN-γ, TNF-α or expression of GZMB, PRF1 among GP33-specific CD8^+^ T cells (Supplementary figure 4f). This suggested that virus-specific CD8^+^ T cells in *Slco^-/-^* are functional upon encountering their cognate antigen, but suffer from impaired activation and expansion.

Once CD8^+^ T cells are activated in secondary lymphoid organs, they migrate into target tissues including the liver. Similar to the results found in the spleen, we found a reduction in LCMV-specific CD8^+^ T cells in the liver (Figure 4c), which was also accompanied by a reduction in total CD8^+^ T cells (Supplementary Figure 5a). There was no difference in the activation state of liver-resident CD8^+^ T cells, however a reduction in total activated cells (Supplementary Figure 5b-c). Although fewer virus-specific T cells were present in the liver of mice with high serum BAs (Figure 4c), it remained unclear whether this would impact their functionality. Interestingly, we observed that once restimulated, CD8^+^ T cells of *Slco^-/-^* mice produced cytokines and cytolytic proteins similar to their littermate controls (Supplementary figure 5d), indicating that CD8^+^ T cells that reach the liver are functional.

Taken together, these results indicate that perturbation in BA metabolism by *Oatp* ablation impairs CD8^+^ T cell activation and expansion in the spleen. As a result, the decrease in virus-specific T cell numbers may impair their ability to infiltrate infected tissues, potentially leading to overall reduced effector functions.

### Loss of BA transporters OATP1A/1B attenuate liver damage

LCMV is a non-cytolytic virus and disease manifestation is primarily driven by immunopathology mediated by CD8^+^ T cells (57). Based on the observed impairment in CD8^+^ T cell activation in the *Slco^-/-^* mouse model, we wanted to assess its impact on pathophysiology and pathogen load upon viral infection.

*Slco^-/-^* mice infected with LCMV exhibited a slightly delayed onset of body weight loss compared to littermate controls (Figure 5a). Based on the reduced virus-specific T cell response, we hypothesized that viral clearance may be affected. Indeed, *Slco^-/-^*mice displayed increased viremia, and higher viral loads in the liver and spleen (Figure 5b). Consistent with the reduced T cell response and increased viremia, we found that genetic ablation of *Slco1a/1b* transporters led to reduced levels of liver damage markers, alanine aminotransferase (ALT) and aspartate aminotransferase (AST) (Figure 5c). Histological assessment of liver pathology showed an increased presence of cellular infiltrates (Figure 5d), which is in line with an increased presence of virus-specific CD8^+^ T cells in the liver of WT mice infected with LCMV Cl13 (Figure 4c). To confirm the heightened inflammatory state of the liver, we also assessed the expression of acute phase proteins and found that these were slightly reduced in *Slco^-/-^* mice (Figure 5e). Together, our data suggest that systemically high levels of BA, which is a feature of several liver diseases including LCMV Cl13 infection, can reduce CD8^+^ T cell activation and thereby prevent excessive CD8^+^ T cell-driven immunopathology.

**Figure 5:**
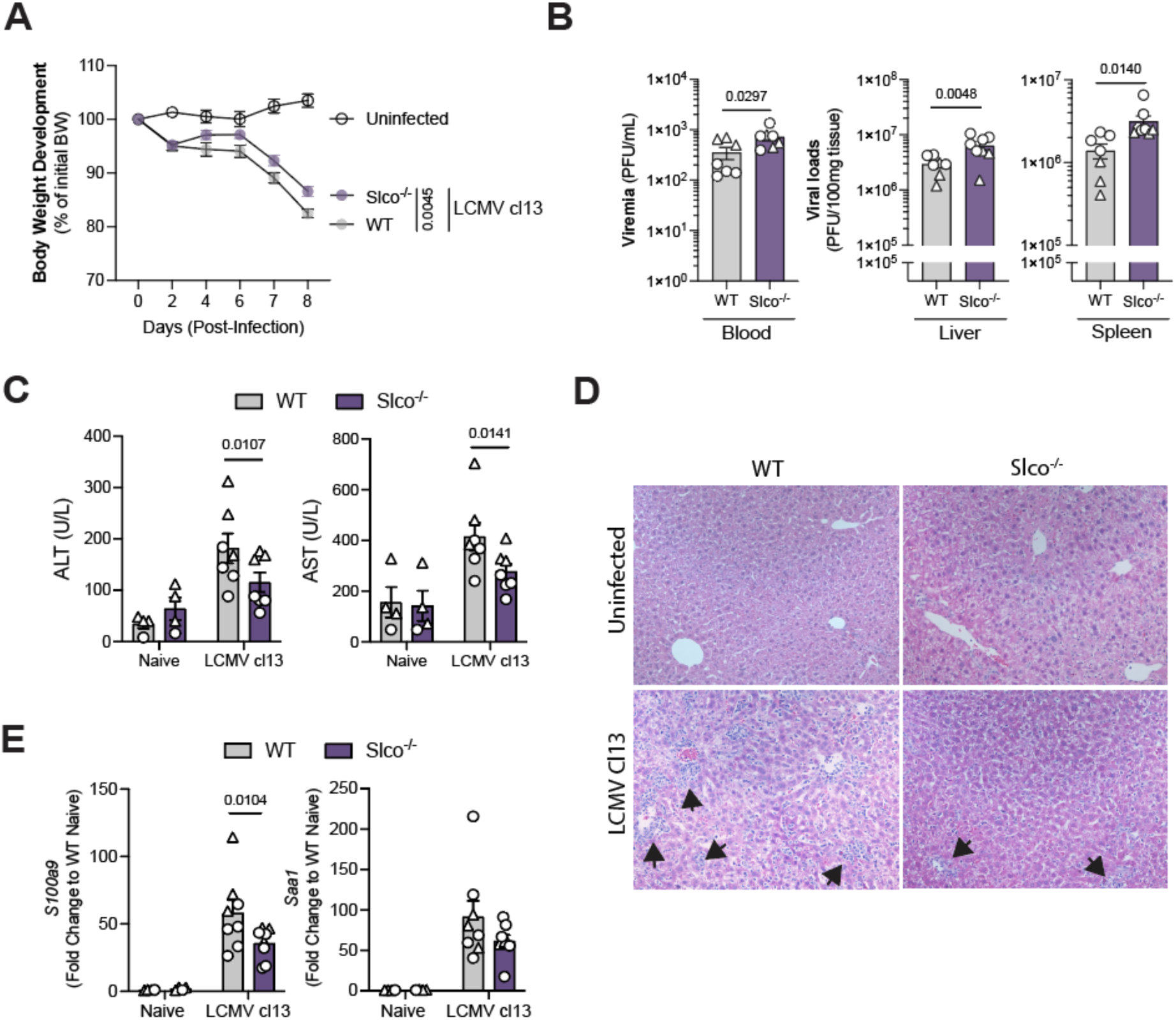
Loss of bile acid transporters *Slco1a/1b* reduces T cell-mediated liver damage during LCMV Cl13. (a) Body weight development of *Slco^-/-^* or littermates infected with LCMV Cl13. (b) LCMV Cl13 titers in blood, liver and spleen at 8 days post-infection in *Slco^-/-^* mice or littermate controls. (c) Serum ALT and AST levels in *Slco^-/-^* mice or littermate controls at 8 days post-infection. (d) Representative H&E staining of liver sections from WT or *Slco^-/-^* at day 8 post infection or left uninfected. Arrows indicate cellular infiltrates. (e) Hepatic gene expression of acute phase proteins in *Slco^-/-^* mice or littermate controls at 8 days post-infection. Data presented as mean ± SEM. Data pooled from two independent experiments; each experiment represented in a different shape (n = 4-8 mice/group). (a,c,e) Two-Way ANOVA. (b) Unpaired Student’s t-test.

## Discussion

In this study, we investigated the interplay of BA metabolism and immunity, demonstrating that CD8^+^ T cell-mediated hepatitis leads to a downregulation of BA-metabolism genes, while also increasing systemic BA levels and shifting their composition. In turn, using mice with a genetic ablation of basolateral OATP transporters as a model of sustained systemically high BA levels resulted in an impaired CD8^+^ T cell response and a concomitant decrease in CD8^+^ T cell mediated immunopathology and viral clearance.

Liver diseases substantially affect systemic BA levels (29,58–61). We found that viral hepatitis induced by LCMV Cl13 increased levels of BAs in the circulation, similarly to LCMV strain WE (47). Our in-depth BA analysis showed that viral hepatitis shifted the BA pool towards liver-abundant host-derived and conjugated BAs, a shift that was also found in patients with chronic hepatitis B infection (27). These similarities establish LCMV as a pathophysiological-relevant mouse model to study BA metabolism.

Additionally, liver diseases, such as cholestatic diseases and viral hepatitis, show a sustained repression of genes involved in BA metabolism (62,63). While the FGF15-FXR-SHP axis is the main driver of this downregulation in cholestatic diseases through the physiological feedback inhibition by BAs in the gut (62), our data indicates that this pathway is not involved in the downregulation of BA transport and synthesis genes in the context of LCMV-induced hepatitis. Instead, the adaptive immune response, specifically CD8^+^ T cells, appear to directly mediate transcriptional downregulation.

Cholestasis is a common feature of inflammatory conditions in the liver (17,18). Basolateral BA export pumps such as MRP3, MRP4, OSTα, and OSTβ also play an important role in cholestatic conditions, allowing hepatocytes to mitigate BA toxicity by eliminating BA back to the blood stream (17) However, our data did not show significant regulation of *Mrp3*, *Mrp4*, *Ostα* or *Ostβ* on day 8 post-infection, suggesting a differential regulation in LCMV-induced hepatitis compared to other inflammatory liver diseases.

Inflammatory cytokines such as IL-6 (64), IL-1β (17) and TNF-α (65) have been shown to control BA transporter expression. Yet, our results suggest that classical pro-inflammatory T cell-derived cytokines, including IL-6, TNF-α and IFN-γ were dispensable for the downregulation of BA transporters, except for *Abcb11*/BSEP, during LCMV-induced hepatitis. Collectively, this highlights the importance of CD8^+^ T cells in regulating BA metabolism either through cytotoxic effects or yet unknown mediators.

Based on our findings, we postulate that the cytotoxic activity of CD8^+^ T cells may be a key driver of viral infection-induced changes in BA metabolism. CD8^+^ T cells activity might be involved in altering BA pools through two distinct, possibly independent mechanisms: (1) release of host-derived BAs from damaged hepatocytes and (2) altered expression of BA transporters responsible for take-up of BAs from circulation. In line with this, Honke *et al.* showed the CD8^+^ T cell dependence of systemic BA elevation in LCMV WE (47). Our findings solidify and establish the role of CD8^+^ cells in driving systemic BA level elevation and the downregulation of hepatic BA metabolism, respectively. Further research will be required to elucidate whether cytotoxic T cells control BA metabolism by directly targeting hepatocytes expressing the BA metabolism machinery.

In a second line of research, we addressed the impact of systemically high BA levels on the immune response. We demonstrate that in a mouse model of systemically high BA levels, CD8^+^ T cell activation is impaired, leading to an associated reduction of viral clearance and CD8^+^ T cell-mediated immunopathology. We propose three plausible mechanisms based on previous findings: certain BA species could directly affect Ca^2+^ signaling (39,66) or mTOR signaling (41) in CD8^+^ T cells. Alternatively, BAs could indirectly modulate T cell responses by affecting dendritic cell cross-presentation (37). Another possible explanation is impaired T cell homing to the liver; however, we observed reduction in virus-specific T cells numbers and their activation marker expression already in the spleen, making this explanation less likely.

Using *Slco^-/-^* mice as a model of high BA level comes with some additional effects including elevated systemic bilirubin, altered energy metabolism and a BA composition distinct from that observed in LCMV infection (11). In addition, some of the genes ablated in *Slco^-/-^* mice have roles in other organs such as the brain and kidney (12), complicating the establishment of clear causal relationships in the *Slco^-/-^* model. Thus, one need to carefully assess how the findings in *Slco^-/-^* mice may translate to LCMV and chronic hepatitis in general.

NTCP/*Slc10a1^-/-^* mice also show profound elevation of (predominantly conjugated) serum BA levels (67), but otherwise lack profound systemic BA effects. In contrast, *Slco^-/-^* mice show and elevation predominantly in unconjugated BA. Individual BA species differ in their ability to penetrate cell membranes and activate BA receptors (68). Thus, further studies should compare our findings with those from mice lacking NTCP/*Slc10a1*, which was beyond the scope of the current study.

In conclusion, LCMV-induced hepatitis shows a perturbed BA metabolism with features akin to liver diseases of various etiologies (6). Intriguingly, the induced alterations in BA metabolism raise the possibility of an immunomodulatory function of BAs to protect from excessive CD8^+^ T cell mediated immunopathology. These findings underscore the importance of systemic BA metabolism in shaping immunity among patients with liver diseases, which represent a global health burden. Using this explorative finding, it is reasonable to investigate the reciprocal interplay between the immune system and bile acid metabolism in liver diseases in greater depth. Such research could reveal novel therapeutic strategies modulating immune responses through metabolic interventions.

## Methods

### Experimental Model and Subject Details

#### Mice

Animals were kept and bred at the Core facility laboratory animal breeding and husbandry of the Medical University of Vienna, under specific pathogen-free (SPF) conditions. Mouse experiments were conducted in individual ventilated cages in compliance with the animal experiment licenses BMWFW-66.009/0361-WF/V/3b/2017 and 2020-0.406.011, approved by the institutional ethical committees of the Department for Biomedical Research of the Medical University of Vienna.

Wild-type mice on C57BL/6J background were bred in-house. *Slco1a1b^-/-^* mice were obtained from Alfred Schinkel (The Netherlands Cancer Institute, Amsterdam, The Netherlands) on an FVB background. These mice were backcrossed onto C57BL/6J for 6 or more generations. For backcrossed mice, we used heterozygous or wild-type littermates as controls. Unless otherwise stated, mice were 8-15 weeks old at the start of experiments, and all mice were age- and sex-matched within experiments. Other genetically modified animals were on C57BL/6 background; *Ifnar1^-/-^* (69) (032045-JAX) and *Rag2^-/-^* (55) (008449-JAX). C57BL/6 males and females were used interchangeably without noticeable difference between readouts.

Pair-feeding experiments were performed on individually housed mice to ensure precise measurement of food intake. They adapted to solitary housing for three days before infection. Infected mice then received food ad libitum, while pair-fed counterparts were given the same exact amount of food consumed by infected mice.

### Infections

Mice were infected intravenously with 2×10^6^ focus forming units (FFU) of Lymphocytic choriomeningitis virus Clone 13 strain (LCMV Cl13). LCMV Cl13 was grown in BHK-21 cells, and viral titers were determined with focus forming assay (70). Mice were sacrificed at the indicated time points, and tissues were snap-frozen in liquid nitrogen to be stored at −80°C until further analysis.

### Treatment with blocking antibodies

For the blocking of cytokines, animals received 0.5 mg/mouse of the following antibodies: anti-IL-6 (MP5-20F3, Rat IgG1, BioXcell #BE0046)), a-IFN-γ (XMG1, Rat IgG1, BioXcell #BE0055), a-TNF-α (XT3.110, Rat IgG1, BioXcell #BE0058) or a rat IgG1 isotype (MOPC-21, BioXcell #BE0083). The injections were given intraperitoneally every other day for seven days, with the first given one day prior to the infection with LCMV Cl13.

For blocking of CD8, animals received 0.2 mg/mouse of anti-CD8 (YTS169.4, Rat IgG2b, BioXcell #BE0117) or a rat IgG2b isotype (LTF-2, BioXcell # BE0090). The injections were given intraperitoneally two, and one day before infection with LCMV Cl13.

### Blood chemistry

For serum analysis, blood samples were centrifuged at 10,000g for 5 minutes at 4°C, after which serum was stored in a new tube at −80°C until further analysis. On the day of the analysis, serum samples were diluted 1:8 in PBS. Alanine aminotransferase (ALT), aspartate aminotransferase (AST), albumin, bilirubin and cholesterol were measured using a Cobas C311 analyzer (Roche). Total BAs in the serum were determined by an enzymatic assay kit (Cell Biolabs #STA-63), according to the manufacturer’s instructions.

### RNA isolation and real-time PCR

The TissueLyser II (Qiagen) was used to homogenize the tissues, and total RNA extraction was performed using Qiazol Lysis Reagent (Qiagen, #79306) following the manufacturer’s instructions. cDNA was synthesized using the RevertAid First Strand cDNA Synthesis Kit (Thermo Fisher Scientific, #K1622). Real-time PCR was performed using a Taqman Fast Universal PCR Mastermix (Thermo Fisher Scientific, #4352042), and Taqman Gene Expression Assays (Thermo Fisher Scientific) against the following mouse gene products: Slco1a1 (#Mm00649796_m1), Slco1b2 (#Mm00451510_m1), Abcb11 (#Mm00445168_m1), Slc10a1 (#Mm00441421_m1), Nr1h4 (#Mm00436425_m1), Cyp7a1 (#Mm00484150_m1), Nr0b2 (#Mm00442278_m1), Saa1 (#Mm00656927_g1), S100a9 (#Mm00656925_m1), Fgf15 (#Mm00433278_m1), Slc10a2 (#Mm00488258_m1). Ef1a was also measured by Taqman chemistry using the following primers:

5’-GCAAAAACGACCCACCAATG-3’, 5’-GGCCTTGGTTCAGGATA-3’, and probe: 5’- [6FAM]CACCTGAGCAGTGAAGCCAG[TAM]-3’.

### Isolation of intrahepatic lymphocytes

Intrahepatic lymphocytes were assessed as previously described (71). In brief, livers were perfused via the portal hepatic vein using cold PBS. The gall bladder was removed and livers were collected in cold PBS. Tissue was mechanically disrupted using a 70µm cell strainer (Sarstedt) to obtain a single-cell suspension. Cells were pelleted at 400g for 5min at 4°C. Cells were then resuspended in a 42% Percoll solution (Cytiva, #17-0891-02) and centrifuged for 20min, 800g at room temperature without break. Red blood cells were lysed by resuspending the pellet in 1 mL of RBC Lysis Buffer (Invitrogen, #00-4333-57) for 3min at room temperature. Cells were pelleted at 400g for 5min at 4°C. Finally, cells were resuspended and used flow cytometry staining.

### Flow cytometry

Spleens were isolated and collected in cold PBS. Single cells were isolated by mechanical disruption of spleens through a 70µm cell strainer (Sarstedt). Cells were spun down at 400g for 5min at 4°C. Pellets were resuspended in 1mL of 1x RBC lysis buffer (Invitrogen, #00-4333-57) for 5min at room temperature. Cells were washed in PBS and spun down again at 400g, 5min at 4°C. Cells were resuspended in cold PBS and used for flow cytometry staining. For PMA/ionomycin restimulation, cells were plated in 96-well plates and then restimulated with Cell Activation Cocktail with Brefeldin A (Biolegend #423303) for 4 hours at 37°C. For viral peptide restimulation, cells were initially incubated with the peptide alone for 30min at a concentration of 1µg/mL at 37°C. Subsequently, at the same peptide concentration, Protein Transport Inhibitor Cocktail (eBioscience, #00-4980-03) was added and cells were incubated at 37°C for 4 hours. After incubation cells were washed and stained for surface markers and intracellular proteins.

For surface staining, cells were incubated with fluorophore-labelled antibodies, Fixable Viability Dye eFluor™ 780 (eBioscience #65-0865-14) and CD16/32 FcR-Block (Biolegend #101302). In case, virus specific tetramers (i.e., PE-labelled GP33-specific tetramers acquired from the NIH Tetramer Core Facility, US), cells were incubated for staining at room temperature for 20min. After cells were washed twice, cells were fixed in 4% PFA for 10min at room temperature. In case of intracellular staining, cells were fixed/permeabilized using the eBioscience™ Foxp3/ Transcription Factor Staining Set (00-5523-00, Invitrogen). Cells were incubated with intracellular staining overnight at 4°C. The following antibodies were used for the staining: TCRb-PerCP-Cy5.5 (Biolegend #109227), CD8a-BV711 (Biolegend #100747), CD44-AF700 (Biolegend #103025), GranzymeB-AF700 (Biolegend #372221), Perforin-PB (Biolegend #154311), TNF-α-APC (Biolegend #506307), IFN-γ-PE-Cy7 (Biolegend #505826), PD1-BV605 (Biolegend #135219), KI67-AF488 (Biolegend #151204).

### Targeted LC-MS based metabolite measurements

Profiles of murine primary and secondary unconjugated and conjugated C24-BAs in serum (30 µl), were analyzed as published previously (72).

### Bioinformatic analysis

Heatmaps were made with the Complexheatmap package (73), and KEGG pathway enrichments were performed using the clusterProfiler package (74) in R version 4.2.2. Relative abundance of bacterial bile salt hydrolase genes (EC:3.5.1.24) in feces of uninfected, LCMV Armstrong or LCMV Cl13 infected mice 8 days post-infection were predicted with picrust2 (75) implemented in qiime2 (76). 16S amplicon sequencing tables with ASVs generated via DADA2 of the Labarta-Bajo *et al*. study (50) have been downloaded from the qiita (77) repository (ID 11043).

### Statistical analysis

The results are presented in the format of mean ± SEM and were subjected to statistical analysis as specified in the figure captions, employing GraphPad Prism. Relevant p-values were presented. Unless otherwise specified in the figure legend, data shows representative results from at least two independent experiments. For pooled data, due to smaller individual experimental group size, each individual experiment was presented with different shapes in the same graph and statistical analysis was blocked for experiment.

## Acknowledgments

We thank Alfred H. Schinkel from The Netherlands Cancer Institute for the provision of *Slco1a1b^-/-^* mice and Irmgard Fischer from the Max Perutz Histology Unit for help with tissue processing.

## Data availability

The metabolomics dataset supporting our findings is available as a supplementary table accompanying this publication.

## Supporting information

Supplementary table: BA profiling

**Supplementary Figure 1:**
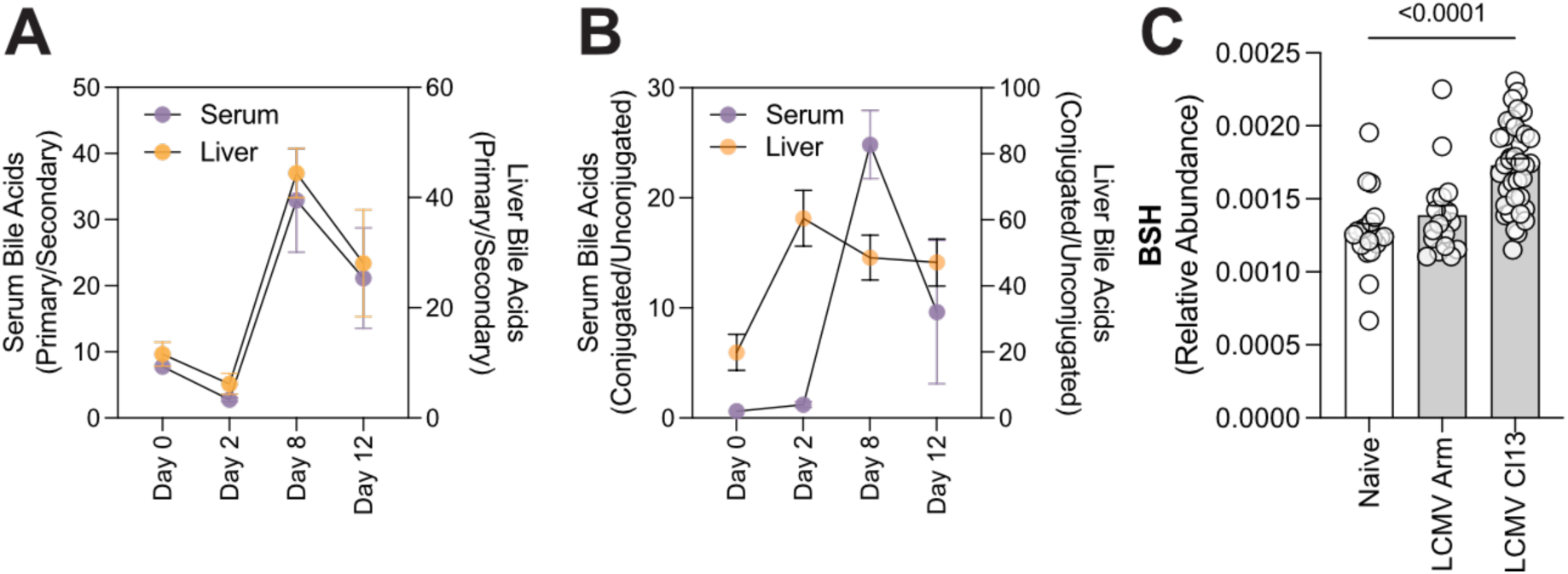
Chronic LCMV infection alters BA composition, presumably independent of BSH activity. (a) Ratio of host-to-microbe-derived BA species in serum and liver over course of LCMV infection. (b) Ratio of conjugated-to-unconjugated BA species in serum and liver over course of LCMV infection. (c) Relative abundance of BSH-expressing microbial species in uninfected, LCMV Armstrong (Arm) or LCMV Cl13 infected mice 8 days post-infection, based on 16S RNA data from Labarta-Bajo *et al*. (50). Data presented as mean ± SEM. Representative data from two independent experiments (n = 4 mice/time point) (a,b). Kruskal Wallis test (c)

**Supplementary Figure 2:**
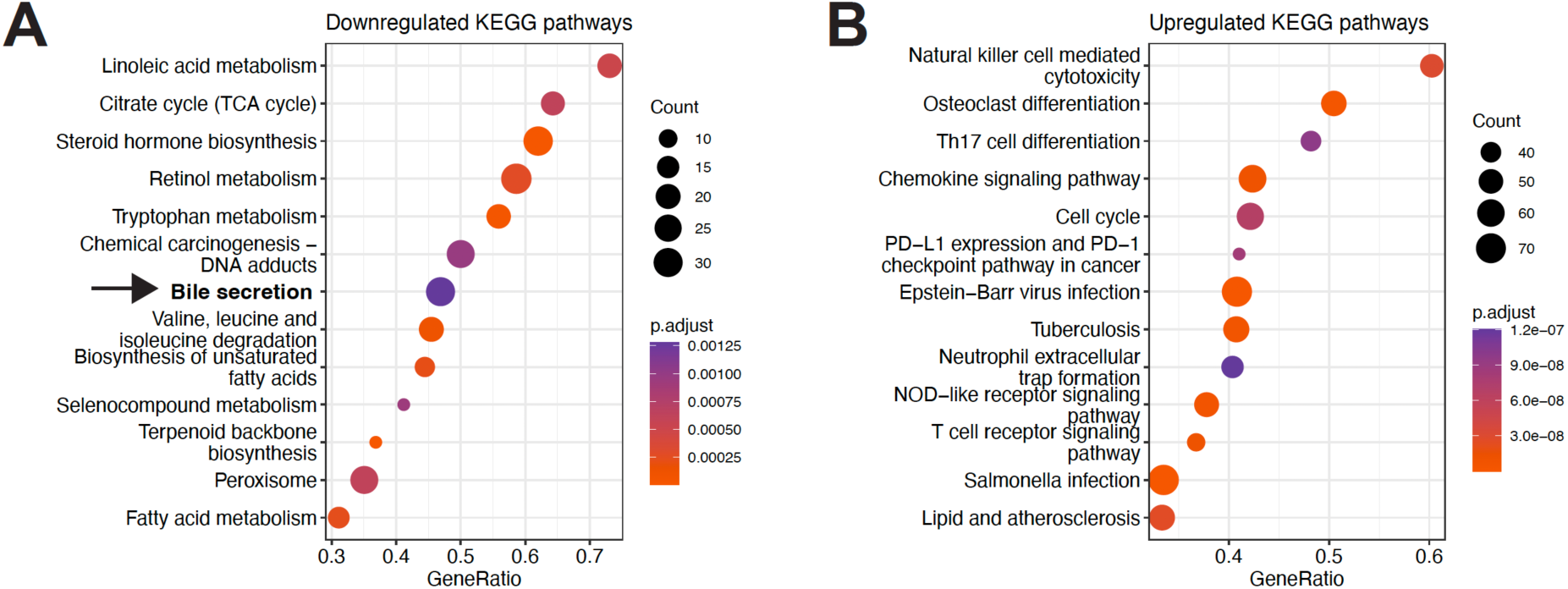
Pathway enrichment analysis of livers in response to LCMV cl13 infection. (a,b) Gene set enrichment analysis of hepatic gene expression dataset (52) at day 8 post-LCMV Cl13 infection for downregulated (a) and upregulated (b) KEGG pathways.

**Supplementary Figure 3:**
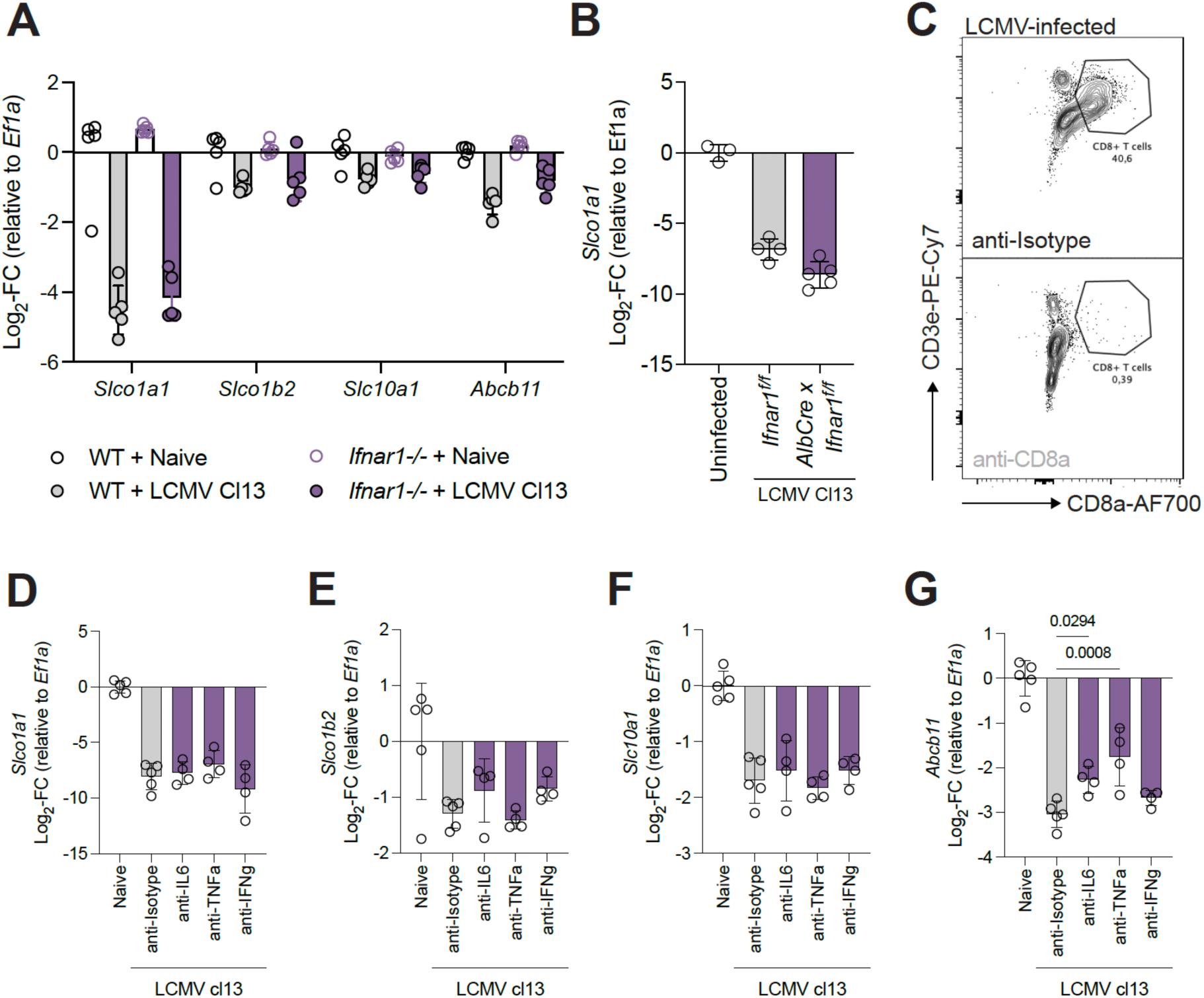
Downregulation of bile acid transporters in the liver is independent of type I signaling and largely independent IL-6, TNF-α and IFN-γ. (a) Expression of BA transporters in *Ifnar*-deficient mice at 8 days post LCMV Cl13 infection. (b) *Slco1a1* expression in liver-specific *Ifnar*-deficient mice at 8 days post LCMV Cl13 infection. (c-f) Expression of hepatic BA receptors in response to LCMV Cl13 and antibody-mediated cytokine blockade at day 8 post infection. Data presented as mean ± SEM. Representative data from one independent experiment (n = 5 mice/group) (a) and from two independent experiment (n = 3-5 mice/group) (b). Representative data from three independent experiments (n = 4-5 mice/group) (c-f). (c-f) One-Way ANOVA.

**Supplementary Figure 4:**
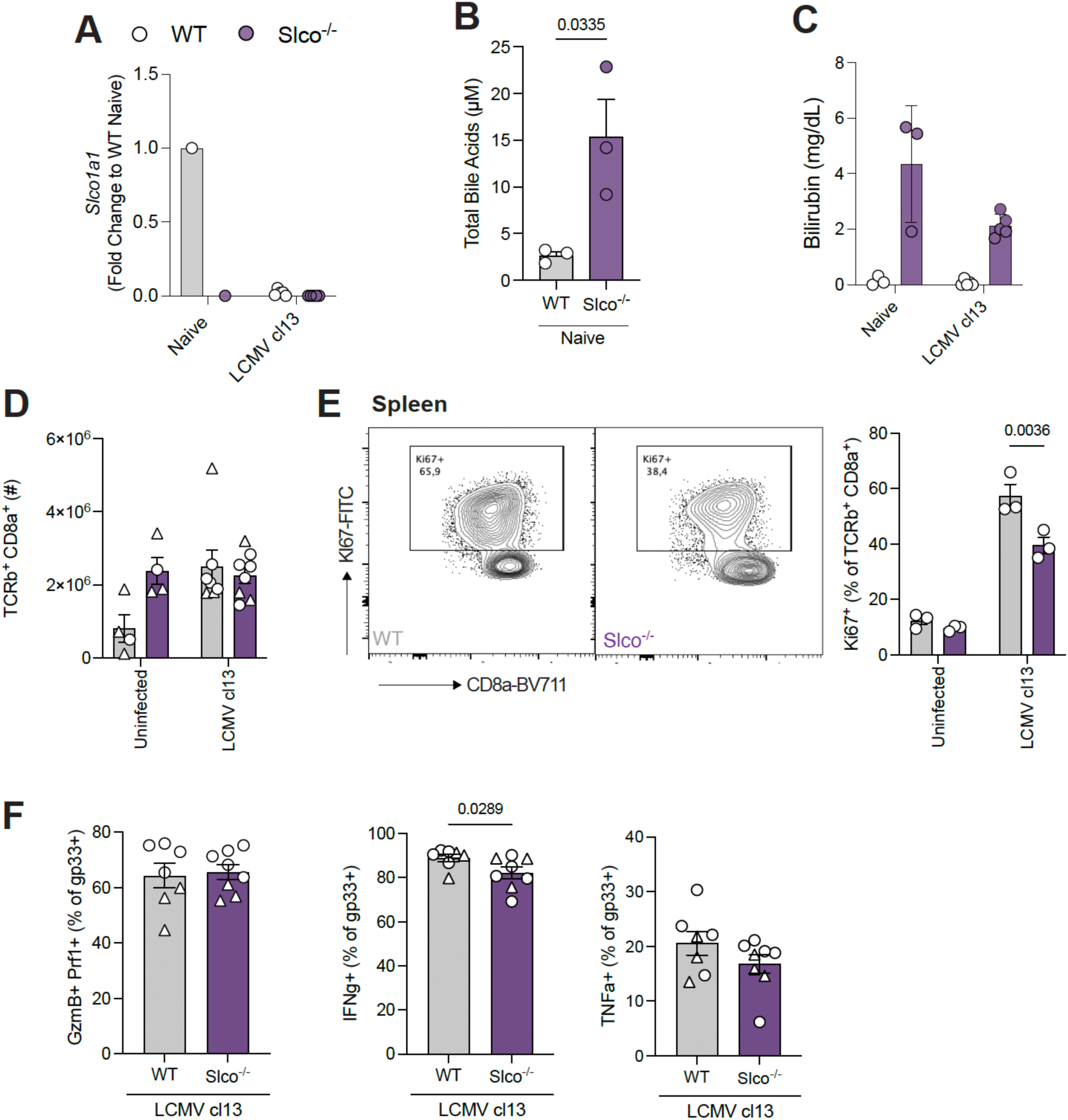
Loss of BA transporter increased total BA and bilirubin levels, and is associated with changes in CD8^+^ T cell numbers and proliferation. (a-c) Assessment of the known effects of *Slco*-deficiency in C57BL6/J background. (a) Expression of *Slco1a1* in the liver in *Slco^-/-^* and littermate controls upon infection with LCMV Cl13. (b) Total serum BA levels in uninfected *Slco^-/-^* and littermate controls. (c) Serum bilirubin levels in *Slco^-/-^* and littermate controls upon infection with LCMV Cl13. (d-e) Total number of CD8^+^ T cells in the spleen (d) of *Slco^-/-^* animals and littermate controls. (e) Representative flow cytometry plot for LCMV KI67-expression of CD8^+^ T cell in the spleen 8 days post LCMV Cl13 infection. (f) Cytokine and cytolytic enzyme expression of CD8^+^ T cells in spleen 8 days post LCMV Cl13 infection. Data presented as mean ± SEM. Representative experiment from one experiment (n = 5 infected mice/group) (a), from one experiment (n = 3 mice/group) (b), from two independent experiments (n = 3-5 mice per group) (c). Data pooled from two independent experiments (n = 4-8 mice/group), each experiment represented in a different shape (d,f). Data from one experiment (n = 3 mice/group) (e). Two-Way ANOVA (d-e). Mann-Whitney test (f).

**Supplementary Figure 5:**
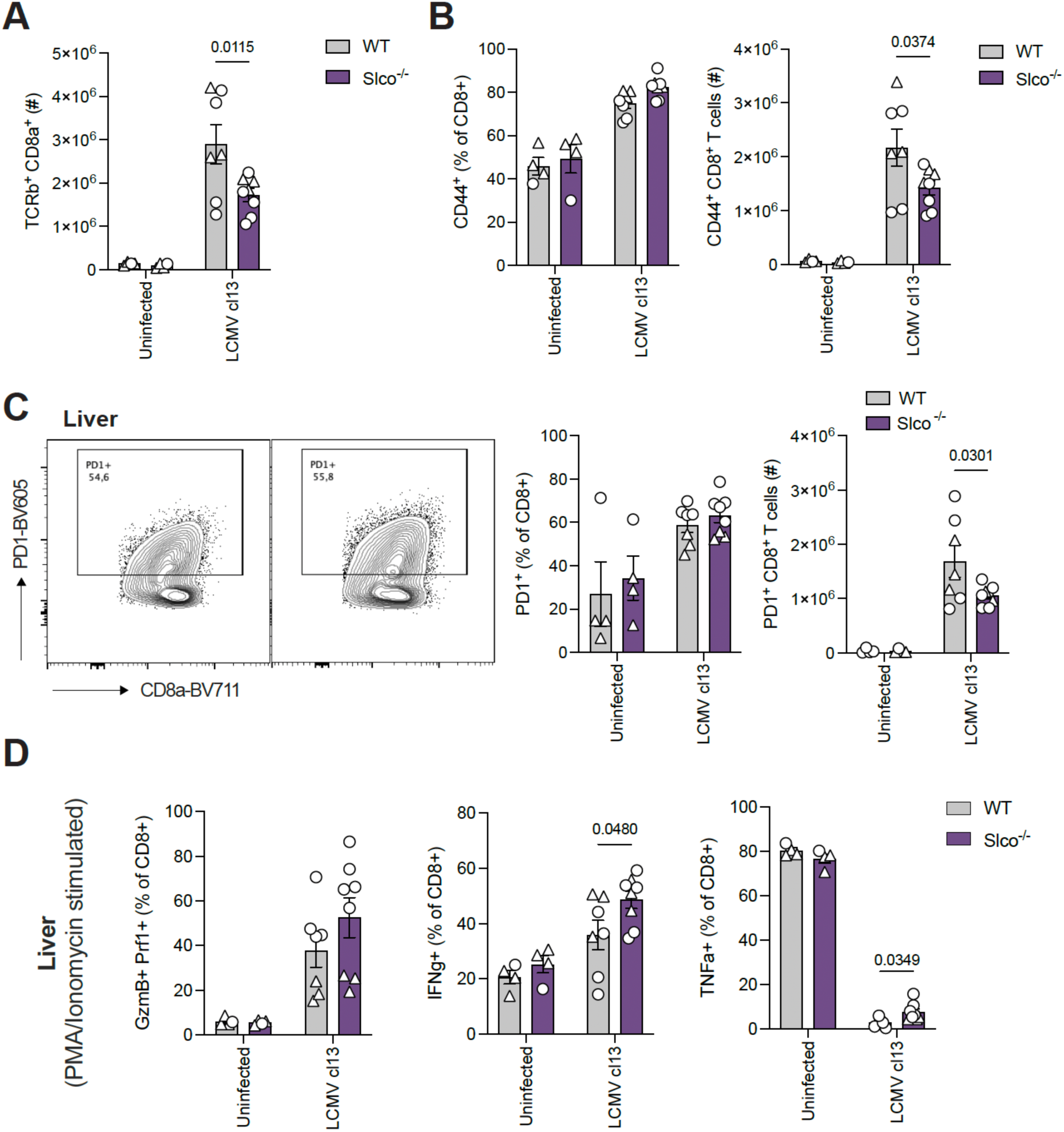
CD8^+^ T cell infiltrating in the liver of *Slco^-/-^* mice is reduced, but virus-specific CD8^+^ T cells are functional. (a) Total number of CD8^+^ T cells in the liver of *Slco^-/-^* animals and littermate controls. (b,c) Frequency of (b) CD44- and (c) PD1-expressing CD8^+^ T cells in liver at day 8 post-LCMV Cl13 infection. (d) Assessment of cytokine production in liver CD8^+^ T cells, isolated at day 8 post LCMV Cl13 infection, upon restimulation with PMA and ionomycin for 4h. Data presented as mean ± SEM. Data pooled from two independent experiments (n = 4-8 mice/group), each experiment represented in a different shape (a-d). Two-Way ANOVA (a-d).

